# Engineering transmembrane signal transduction in synthetic membranes using two-component systems

**DOI:** 10.1101/2022.10.30.514420

**Authors:** Justin A. Peruzzi, Nina R. Galvez, Neha P. Kamat

## Abstract

Cells use signal transduction across their membranes to sense and respond to a wide array of chemical and physical signals. Creating synthetic systems which can harness cellular signaling modalities promises to provide a powerful platform for biosensing and therapeutic applications. As a first step towards this goal, we investigated how bacterial two-component systems can be leveraged to enable transmembrane-signaling with synthetic membranes. Specifically, we demonstrate that a bacterial two-component nitrate-sensing system (NarX-NarL) can be reproduced outside of a cell using synthetic membranes and cell-free protein expression systems. We find that performance and sensitivity of the two-component system can be tuned by altering the biophysical properties of the membrane in which the histidine kinase (NarX) is integrated. Through protein engineering efforts, we modify the sensing domain of NarX to generate sensors capable of detecting an array of ligands. Finally, we demonstrate that these systems can sense ligands in relevant sample environments. By leveraging membrane and protein design, this work helps reveal how transmembrane sensing can be recapitulated outside of the cell, adding to the arsenal of deployable cell-free systems primed for real world biosensing.

**Significance Statement:** Cells detect and respond to environmental and chemical information by using a combination of membrane proteins and genetic polymers. Recapitulation of this behavior in synthetic systems holds promise for engineering biosensors and therapeutics. Using the nitrate-sensing bacterial two-component system as a model, we demonstrate methods to reproduce and tune transmembrane signaling in synthetic lipid membranes, leading to the synthesis of genetically programmed proteins. Through this study, we gain insight into how membrane augmented cell-free systems can be used as a platform to characterize membrane-receptor interactions and engineer new biosensors.

## Introduction

Cell-free transcription and translation systems are powerful tools to study and engineer biology (*1*). Initially used to decipher the genetic code (*2*), cell-free systems have now been used for applications ranging from the development of therapeutics (*3–5*) to distributable biosensors (*6–8*). Recently, synthetic membranes have further expanded the capabilities of cell-free systems, most often acting as a compartment to concentrate and protect encapsulated components (*9–13*). Yet increasingly, membranes have been recognized for their capacity to augment cell-free systems by incorporating functional transmembrane proteins (*9*), enabling protein posttranslational modification (*14*), lipid biosynthesis (*15*), and membrane permeability (*16, 17*). A critical gap in the design of membrane-augmented cell-free systems to date, however, has been the integration of transmembrane signaling capabilities that both leverage the diverse array of membrane proteins and engage with genetic systems. To date, cell-free systems used with synthetic membranes have largely been designed to detect membrane permeable molecules or physical cues, such as light, prior to engaging gene expression systems (*11, 13, 18, 19*). While this approach has greatly expanded the capabilities of cell-mimetic systems, it is limited by the narrow number of analytes and signals that can be sensed, ultimately restricting the application of cell-mimetic technologies. Transmembrane receptors that can transduce a wide array of chemical and environmental signals across membranes into genetically programmed responses will greatly expand the functionality of cell-free systems, enabling biosensing and programmed biosynthesis in complex aqueous environments, such as the body.

Widespread in prokaryotes, two component systems (TCS) are simple transmembrane sensing motifs which enable the transduction of environmental stimuli into a cellular response (*20*). The canonical TCS is composed of a transmembrane protein sensor histidine kinase and a soluble response regulator which can regulate gene expression upon phosphorylation. These systems can sense many unique physical and chemical signals, such as light, osmotic pressure, small molecules, and ions, and can trigger cellular responses ranging from cell division to taxis (*20–23*). Further, TCSs have been shown to be amenable to protein engineering due to their structurally conserved and modular parts (*24–27*), and functional in non-native organisms (*28, 29*). Combined, these features make membrane bound TCSs an attractive system to integrate into cell-free systems.

Here, we investigate how a model membrane-bound TCS, the nitrate sensing NarX-L, can be integrated into synthetic membranes using a cell-free protein synthesis system. We demonstrate that NarX-L can be reconstituted into synthetic membranes and characterize how membrane physiochemical properties can be used to tune the activity of the sensor. Further, we use protein engineering to explore the modularity of membrane-bound kinases to sense other ligands, and we find that the activity of the histidine kinase depends on both the membrane composition and sensing domain. Using this system, we generate nanoparticles that can sense multiple ligands, detect contaminants in nonideal matrices, and remain functional when fully encapsulated into synthetic vesicles.

## Results

### Integration of the two-component system, NarX-L, into synthetic membranes

As a first step towards understanding how TCSs can be integrated into cell-free systems, we tested the functional expression of NarX and NarL. NarX-L was chosen because the analyte for this sensor, nitrate, is a known ground water contaminate (*30*), serves as a convenient input, and previous work has characterized and engineered NarL mutants and its cognate promoter (*26*), ensuring that downstream gene transcription is a result of cell-free expressed NarL. We assembled cell-free reactions containing the myTXTL system (*31*), three DNA templates, liposomes, and assessed the reactions in response to nitrate addition. Two templates contained genes for either NarX or a NarL mutant (*26*) under the control of T7 RNA polymerase. The third template served as a genetic reporter for TCS signaling and contained a reporter gene (nanoluciferase) under the control of a promoter to which NarL binds (*32*). Finally, we included 100 nm lipid vesicles composed of 1,2-dimyristoyl-sn-glycero-3-phosphocholine (DMPC) into which NarX could integrate (Fig. 1A). Upon addition of nitrate, NarX should phosphorylate NarL, which should then bind the reporter plasmid and induce downstream luciferase expression (Fig. 1A).

**Figure 1.**
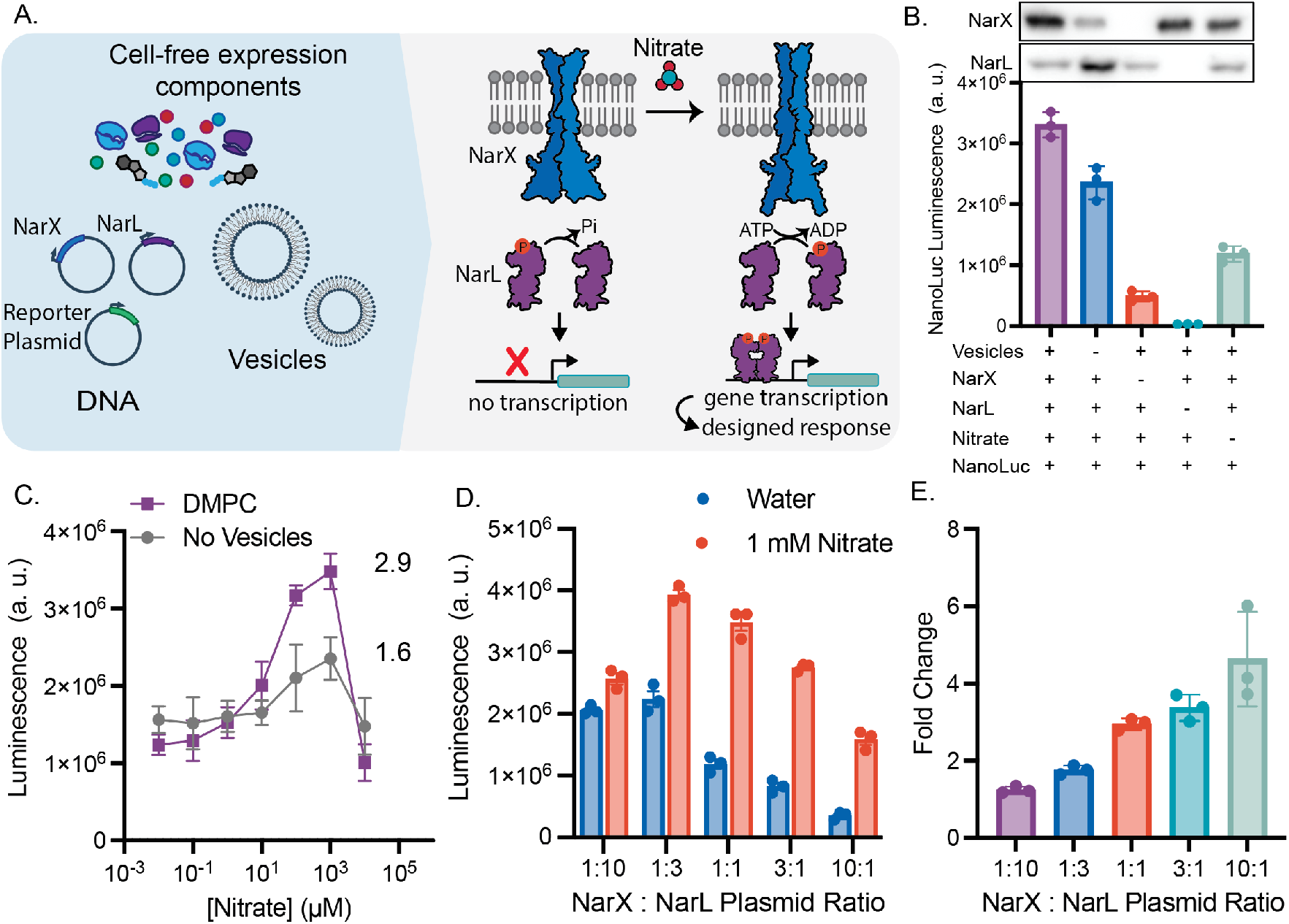
Model two component system NarX-L is readily integrated and active in synthetic membranes. (A) Cell-free reactions were assembled by combining proteins required for gene transcription and translation, plasmids encoding T7 RNAP, NarX, NarL, a reporter gene (nanoluciferase), a membrane mimetic (DMPC liposomes), and nitrate. Upon expression and binding to nitrate, NarX can phosphorylate NarL. Phosphorylated NarL then dimerizes, binds to the promoter, and initiates transcription of the reporter gene, nanoluciferase. (B) The highest luminescence is achieved when all components of the sensor are present. Components were systematically removed to characterize downstream luciferase expression. (C) Cell-free expressed NarX-L has higher nitrate induced expression of luciferase in the presence of a membrane mimetic, DMPC. (D) Sensor activity as reported by luminescence and (E) fold change of NarX-L can be tuned by altering the DNA ratio of NarX and NarL. By increasing the DNA ratio of NarX : NarL, the overall luminescence signal is decreased, but the fold change in luminesce in response to nitrate is increased. The sum of NarX and NarL plasmid concentrations were kept at 6.6 nM. All error bars represent the S. E. M. for *n=3* independent replicates.

We first tested NarX-L function in our cell-free system by monitoring NarX and NarL expression and luciferase luminescence as a function of reaction components. We observed the highest luminescence when all components were present, about a 3-fold increase compared to reactions run in the absence of nitrate, indicating that nanoluciferase expression depends on both nitrate (Fig. 1B) *and* NarX and NarL expression. We confirmed NarX integration into vesicle membranes via immunoprecipitation (Fig. S1). When vesicles were removed from the reaction, we still observed luminescence, likely due to nonspecific binding of NarL to its cognate promoter and subsequent transcription when NarL is expressed at elevated levels. This non-specific interaction of NarL with the reporter plasmid is further supported by the dramatic reduction in luminescence upon removal of the NarL plasmid (Fig. 1B).

We then evaluated nitrate sensitivity and the role of membranes in our system by conducting a titration of nitrate with reactions containing either no vesicles or DMPC vesicles. In the absence of vesicles, we observed a 2-fold increase in luminescence indicating either that NarX may be cotranslationally inserting into native vesicles present in the extracts (*33, 34*) or there may be active, native NarX present in lysates (*35*). When DMPC vesicles were present, we observed a higher maximum luminescence, suggesting that NarX activity is enhanced by the synthetic vesicles, likely through cotranslational integration into the vesicle membranes (Fig. 1C). We determined our cell-free system allowed for detection of nitrate at levels as low as 10 µM, which is below the EPA limit (~100 µM)(*30*). Our cell-free system lost sensitivity when nitrate concentrations were increased to 10 mM nitrate, as higher levels of nitrate inhibit protein synthesis (Fig. S2).

We then sought to tune sensor performance by altering the expression of NarX and NarL. In cell free systems, protein expression can be easily tuned by altering the amount of plasmid present in reactions (Fig. S4) (*36*). By retaining the total amount of NarX and NarL plasmid (6.6 nM), but altering the ratio of NarX and NarL plasmid from 1:10 to 10:1, we observed how expression impacted the expression of luciferase and the sensitivity of the sensor (Fig. 1 D, E). Increasing the amount of NarL plasmid increased luminescence in the absence of nitrate, likely due to nonspecific binding of NarL to the promoter (Fig. 1 D). When NarX plasmid concentration was increased relative to NarL, we observed a reduction in the overall signal but an increase in fold change, calculated as the ratio of luminescence in the presence of 1 mM nitrate to luminescence in the absence of nitrate (Fig. 1 E, S3). These experiments demonstrate that the NarX-L can be functionally expressed in cell-free systems, and its performance can be tuned by altering the ratio of NarX and NarL plasmid.

### Modulation of NarX activity through the tuning of membrane biophysical features

We next wondered if we could further tune sensor performance by modulating biophysical properties of the membranes present in the cell-free reaction. Membrane physical features, (such as lipid chain length and saturation, charge, curvature, and fluidity) have been shown to alter protein activity (*37–40*), as well as the cotranslational insertion and folding of membrane proteins (*16, 41–43*). To investigate the role of chain saturation on NarX-L performance, we assembled cell-free reactions with membranes composed of DMPC (14:0), POPC (18:1-16:0), and DOPC (18:1). These compositions should yield bilayer vesicles with membranes that are composed of lipids with two saturated acyl chains, one saturated and one unsaturated acyl chain, and two unsaturated chains, respectively. We further tested NarX-L activity in membrane systems with 30% POPE to investigate the impact of negative curvature lipid, 30% POPG to investigate the impact of membrane charge, and in 100% *E. coli* polar lipid extract, NarX’s native membrane environment (Fig. 2A). We assembled cell-free reactions with 100 nm vesicles containing these different membrane compositions with and without 1 mM nitrate. We found that using POPC membranes in cell-free reactions yielded the highest luminescence in the presence of nitrate and induced the highest fold-change in response to the addition of nitrate (Fig. 2B).

**Figure 2.**
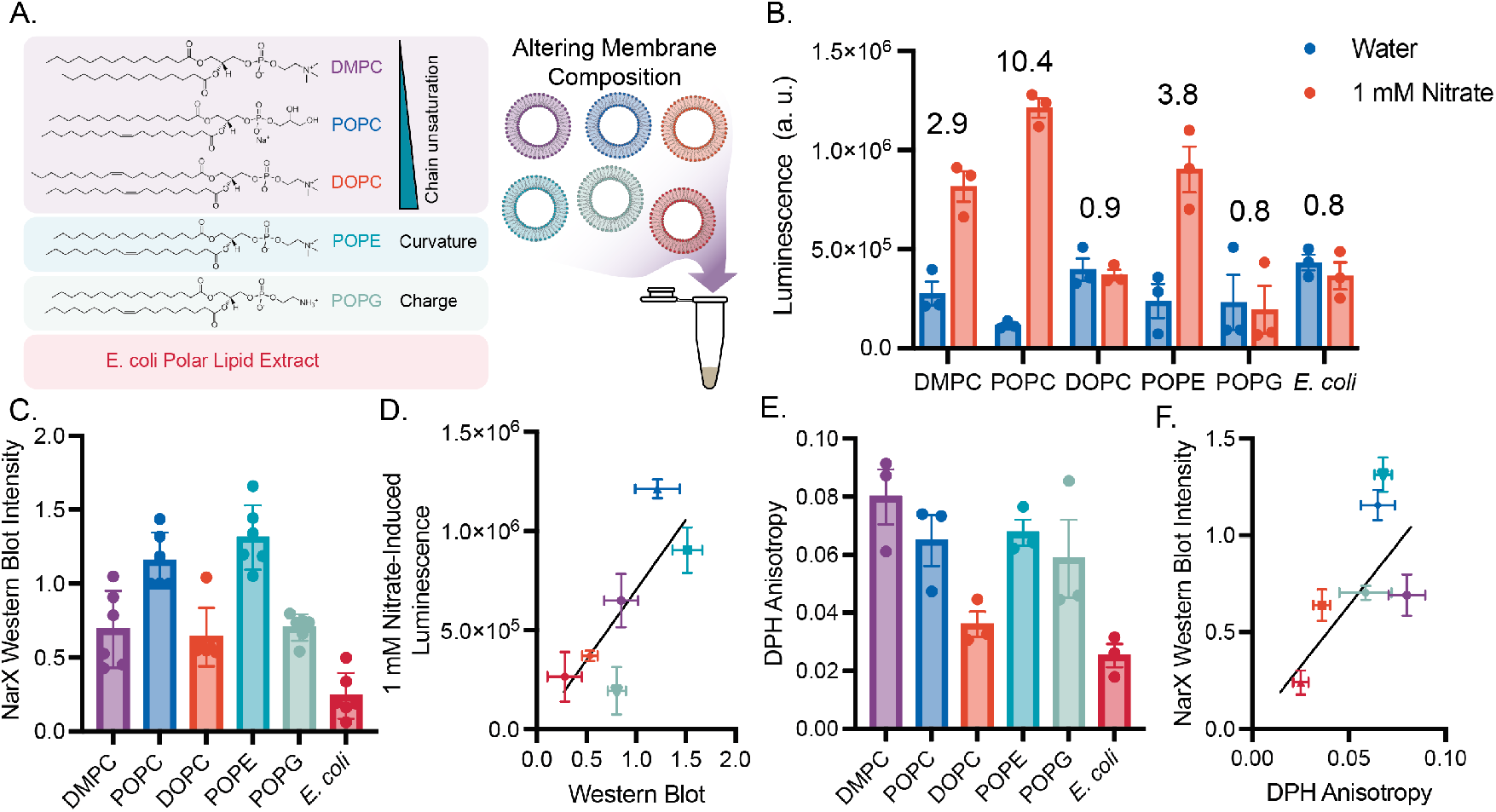
Vesicles of varying composition alter NarX and NarL expression and subsequent activity. (A) By modulating the composition of vesicles that are doped into cell-free reactions, we can characterize how membrane physical features affect NarX cotranslational insertion and activity. (B) NarX-L activity in membranes of different composition in the absence and presence of 1 mM nitrate. Fold change in luminescence is indicated over each membrane composition. (C) NarX expression varies with membrane composition as determined by western blot band intensity. (D) Western blot band intensity is correlated with luminescence and performance of the sensor. (E, F) The viscosity of each membrane measured via DPH anisotropy correlates with protein expression as determined by western blot. All error bars represent the S. E. M. for *n=3* independent replicates. For western blots, *n=6* as band intensities in the absence and presence of nitrate were grouped for each membrane condition.

We wondered if membrane composition impacted protein expression and thereby affected the cell-free sensor performance. To better understand how membranes affect protein expression, we performed western blots on our reactions to quantify NarX and NarL expression (Fig. 2C). We found that NarX expression is correlated with NarL and luciferase expression. We suspect that expression and proper folding of NarX likely enhances the yield of NarL and the downstream nanoluciferase expression in cell-free reactions (Fig. 2D, S5). Further, improper membrane protein folding has been found to increase aggregation and truncation products (*41*), and likely inhibits efficient expression of all protein components in the system. As a result of this relationship, we only considered NarX expression in our future analysis of the effect of membrane composition on NarX-L performance.

To better understand what properties of the membrane enable efficient protein expression, we measured membrane fluidity via DPH fluorescence anisotropy and lipid packing via Laurdan generalized polarization (GP) and compiled other lipid physical properties reported in literature (Fig. 2E, S6, Table S1). Lipid properties are interrelated and often non-monotonic, thus making it difficult to directly relate a single membrane property to protein insertion and activity. To gain a picture of what properties may affect protein yield, we calculated Pearson’s and Spearman’s coefficients for all compiled membrane measurements to protein activity and expression (Fig. S7). Interestingly, we found that DPH anisotropy, which is used as a measure of membrane viscosity, positively correlated with NarX expression (Fig. 2E). Additionally, membrane lateral pressure, lipid neutral plane distance, and membrane transition temperature values compiled from literature also correlated with NarX expression. These features are affected by the structure of the hydrophobic region of the membrane and greatly affect membrane viscosity, further suggesting that membrane viscosity is a key feature for proper cotranslational insertion of NarX (Fig. S7).

To investigate how membrane physiochemical interactions affect NarX activity, we made POPC vesicles and doped 10 mol% of an additional lipid into the membrane (Fig. 3A). POPC was chosen as the main membrane component as it was found to yield the largest fold change in luminescence upon the addition of nitrate (Fig. 2B). By only varying 10 mol% of the lipids in a given membrane composition and keeping the other 90 mol% consistent, we could ensure protein expression of NarX (and correspondingly NarL) remained constant across membrane compositions, thus allowing activity to be attributed to differences in protein conformational changes due to an altered membrane environment instead of total NarX-L protein present in each reaction (Fig. 3B). We again investigated the effect of chain saturation (DMPC, POPC, DOPC), charge (POPG), curvature (POPE), and *E. coli* native membrane lipids on NarX-L activity. POPC still yielded the best fold-induced luminescence, however, doping POPG into membranes yielded the largest increase in luminescence. Further, POPE yielded high luminescence in the presence of nitrate, but interestingly we observed an increase in luminescence in the absence of nitrate (Fig. 3C). This nitrate-independent increase in luminescence upon addition of POPE may suggest that negative lipid curvature stabilizes the “on” conformation of NarX (*44*). The high induction of luciferase expression in POPE and POPG membranes is unsurprising as PE and PG make up the majority of lipid headgroups in *E. coli* membranes (*45*). Adding *E. coli* polar lipid extract yielded a fold change in luminescence close to that of POPG, but a lower luminescence. DOPC still yielded a fold change less than 1, suggesting that increased fluidity may not support proper conformational changes required for induced phosphorylation of NarL and downstream expression upon binding to nitrate. To better understand what features of membranes may enhance sensor performance, we again performed Pearson and Spearman’s correlations (Fig. S8). Physical parameters such as critical packing parameter (CPP), which describes the geometric shape of a lipid, and membrane thickness, as well as membrane mechanical properties, such as bending rigidity (K_c_), were found to correlate with sensor performance (Fig. 3D, E, Table S1). Together, these experiments demonstrate that NarX-L expression and sensitivity can be tuned by altering the physiochemical properties of the membrane mimetic present in the cell-free reaction.

**Figure 3.**
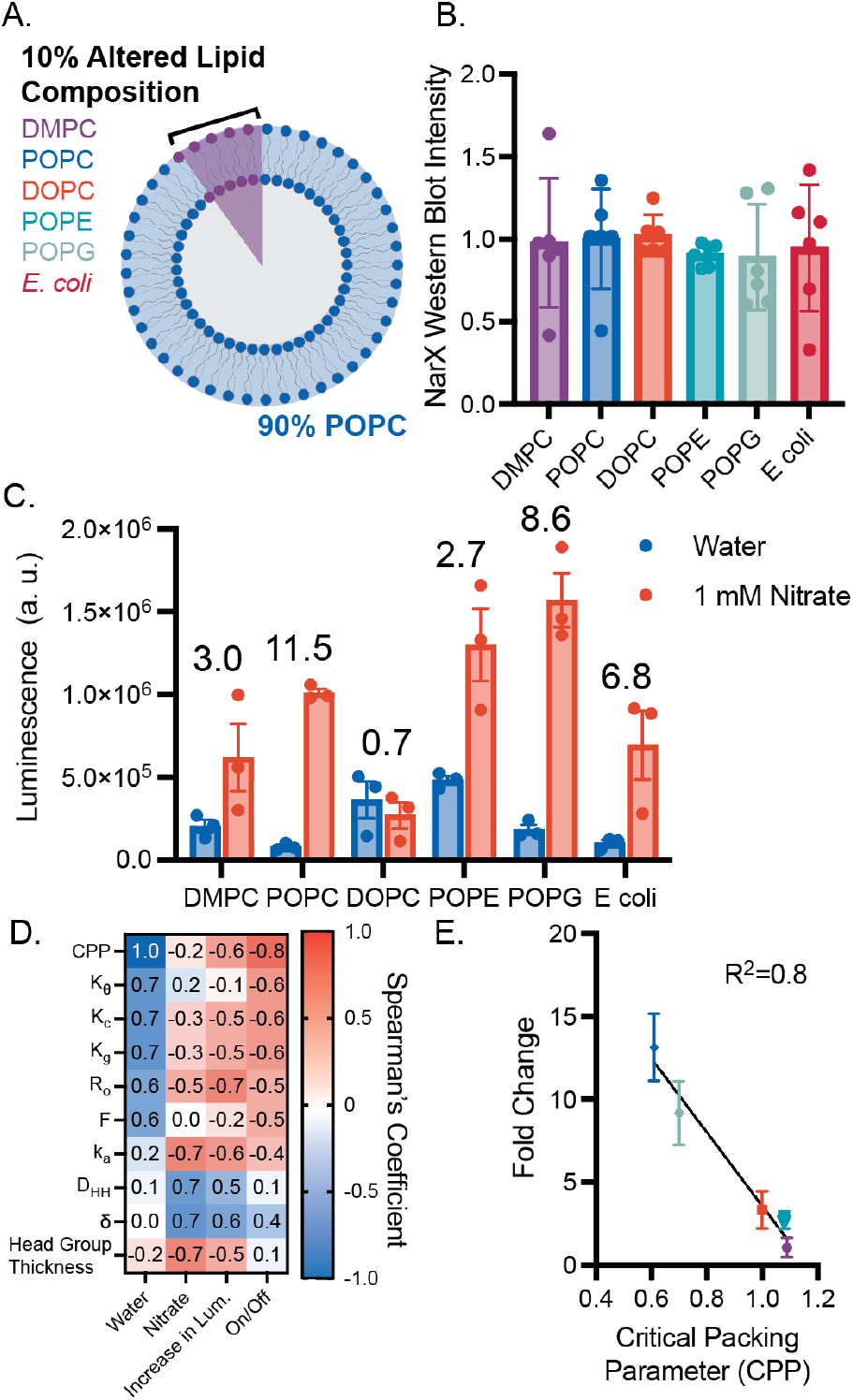
Minor augmentations in membrane composition enable interrogation of lipid physical properties on NarX activity. (A) By altering 10 mol% of the lipid in vesicles composed of POPC, protein expression can be conserved and allow differences in activity to be attributed to protein-lipid interactions. (B) Western blot band intensity of NarX for all samples shows that protein expression was not substantially different when membrane composition was changed. (C) NarX activity and fold change in luminescence varied with membrane composition. (D) Spearman’s correlation demonstrates that changes in luminescence and on/off ratio correlate with lipid physical properties. (E) Lipid critical packing parameter was found to linearly correlate with the fold change in luminescence. All error bars represent the S. E. M. for *n=3* independent replicates.

### Expanding the ligands that can be sensed by the membrane bound histidine kinase NarX

Once we established that plasmid concentrations and membrane physiochemical properties could be leveraged to tune sensor function, we then wondered if we could generate new biosensors via protein engineering. Two-component systems are quite modular and amenable to protein engineering for the design of entirely new sensing and signaling systems in bacterial (*24–26, 46*) and mammalian systems (*28*). We therefore wondered if we could swap out the transmembrane and ligand binding domains of NarX, to generate entirely new cell-free sensors. To accomplish this, we performed a sequence alignment on a subset of membrane bound histidine kinases, and selected candidates based on their alignment to NarX’s HAMP domain, a helical region critical for transmitting ligand binding to kinase activity (Fig. 4A) (*21, 46*). From this analysis, we engineered copper, nickel, iron, and vancomycin sensing NarX chimeras. Specifically, we replaced the transmembrane and sensing domains, as well as the first half of the HAMP domain, of NarX with those from other proteins (NrsS, RssA, VanS, CusS), but retained the histidine kinase (Fig. 4B). By swapping out the transmembrane and sensing domains, but retaining the kinase, we could leverage the same transcription factor and promoter to control downstream gene transcription in response to new ligands.

**Figure 4.**
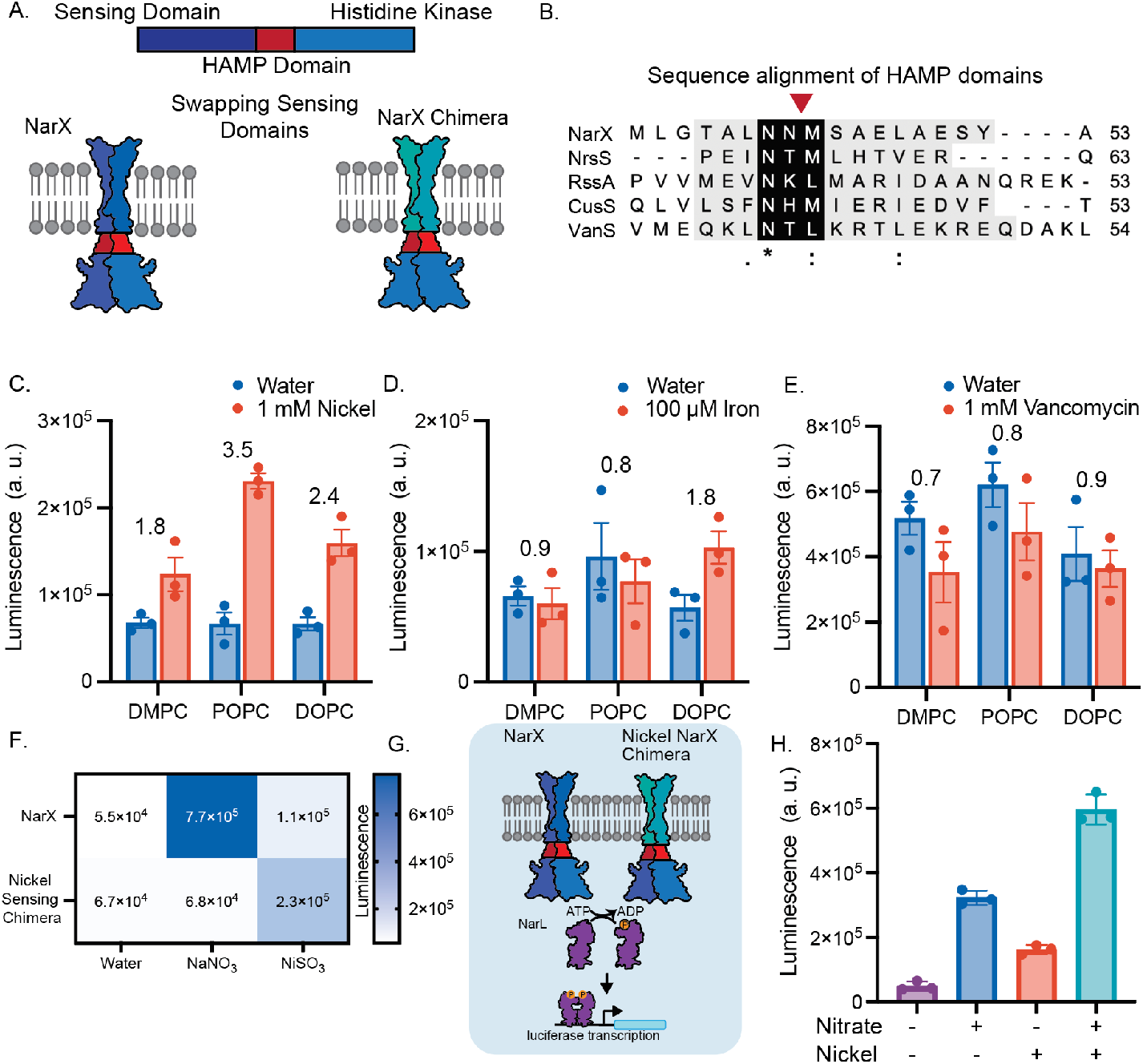
Engineering new NarX chimeras for expanded sensing capabilities. (A) Membrane bound histidine kinases generally possess similar structural features, allowing chimeric proteins with new functions to be engineered by mixing and matching parts. (B) Sequence alignment of NarX (nitrate), NrsS (nickel), RssA (iron), and VanS (vancomycin) HAMP domains. Asterisk (*), colon (:), and periods (.) indicate fully conserved amino acids, or amino acids with strongly conserved, and weakly conserved properties respectively. The red arrow indicates the cross over point of chimeras. (C, D, E) Luminescence as a result of NrsS, RssA, and VanS chimera signaling when expressed in POPC, DOPC, or DMPC membranes with each chimera’s respective ligand. Fold change in luminescence in response to each ligand is indicated above each membrane composition. (F) NarX and NrsS activate specifically to their respective ligands. (G, H) Coexpressing NarX and the NrsS chimera allows for sensing of nitrate and nickel in POPC vesicles. All error bars represent the S. E. M. for *n=3* independent replicates.

We identified several sensing domains that were active in response to the ligand of interest. Kinase and nanoluciferase expression were able to proceed upon addition of the nickel-, iron-, and vancomycin-sensing NarX chimera templates and their ligands. Unfortunately, the addition of copper inhibited cell-free protein expression (Fig. S9). Using plasmid concentrations, which enabled functional NarX sensing, yielded a functional nickel sensor, but no gene expression was observed for the iron and vancomycin sensors in response to each corresponding ligand. We tuned down the concentration of the reporter plasmid by 90% to reduce background expression of nanoluciferase and enhance the specificity of the sensors (Fig. S10). We then expressed proteins in the presence of DMPC, POPC, and DOPC vesicles (Fig. 4C, D, E). Upon reduction of reporter plasmid, we generated functional sensors in response to nickel and iron. Interestingly, each sensor performed differently in each membrane, suggesting that each protein requires specific membrane physiochemical properties for proper function. The vancomycin sensor did not induce an increase in luciferase expression in any of the membrane compositions tested in response to vancomycin. However, luminescence for the vancomycin sensor was highest across all conditions (Fig. 4E), suggesting that the chimera may be locked into an “on” conformation, perhaps due to destabilizing the HAMP domain (*44*). These results demonstrate how protein and membrane engineering can be used to readily generate new cell-free receptors by tapping into the large repertoire of known sensing domains of transmembrane proteins, and underscores the important relationship between membrane physical properties, protein sequence, and function.

Once we established that we could successfully sense multiple ligands, we wondered if we could co-express multiple kinases into vesicles to sense more than one ligand at a time. To test this, we chose to co-express NarX and the nickel-sensing chimera as they both exhibit activity in POPC membranes. We first established the ability of wild-type NarX and the nickel-sensing chimera to specifically turn ‘on’ in response to nitrate and nickel (Fig. 4F). Once specificity was confirmed, we co-expressed wild-type NarX and the nickel-sensing chimera into POPC vesicles in the presence of water, 1 mM nickel, 1 mM nitrate, or 1 mM nickel and 1 mM nitrate (Fig. 4G). We found that upon co-expression, we could sense multiple ligands. Interestingly, we observed the highest luminescence in the presence of both ligands, perhaps because both kinases were able to initiate transcription, leading to increased luciferase expression (Fig. 4H). Integrating multiple receptors into synthetic membranes together with the design of distinct genetic reporters could lead to synthetic vesicles capable of multiplexed sensing and response to many diverse stimuli.

### Priming cell-free expressed two-component systems for real world biosensing

Over the last decade, many promising cell-free systems have been developed, enabling the delivery and sensing of a wide array of ligands in non-ideal environments. Cell-free systems have been encapsulated and administered to mice (*47*), and lyophilized and used in resource limited settings (*6, 7*), and integrated into wearable materials to sense environmental contaminants (*8*). Towards the goal of creating cell-mimetic particles which can sense environmental stimuli in complex environments, such as the body, we encapsulated our cell-free system into giant unilamellar vesicles using the emulsion transfer method (Fig. 5A). We encapsulated cell-free extract containing plasmids encoding NarX, NarL, and the downstream protein (luciferase or mEGFP) into membranes composed of 1 POPC: 1 cholesterol and a rhodamine conjugated lipid. Vesicles encapsulating the reactions were incubated at 30°C for 16 hours with RNase, to prevent transcription outside of the vesicles, and 1 mM nitrate. We observed that the sensor remained functional and was able to sense exogenous nitrate via microscopy and a bulk plate reader assay (Fig. 5B).

**Figure 5.**
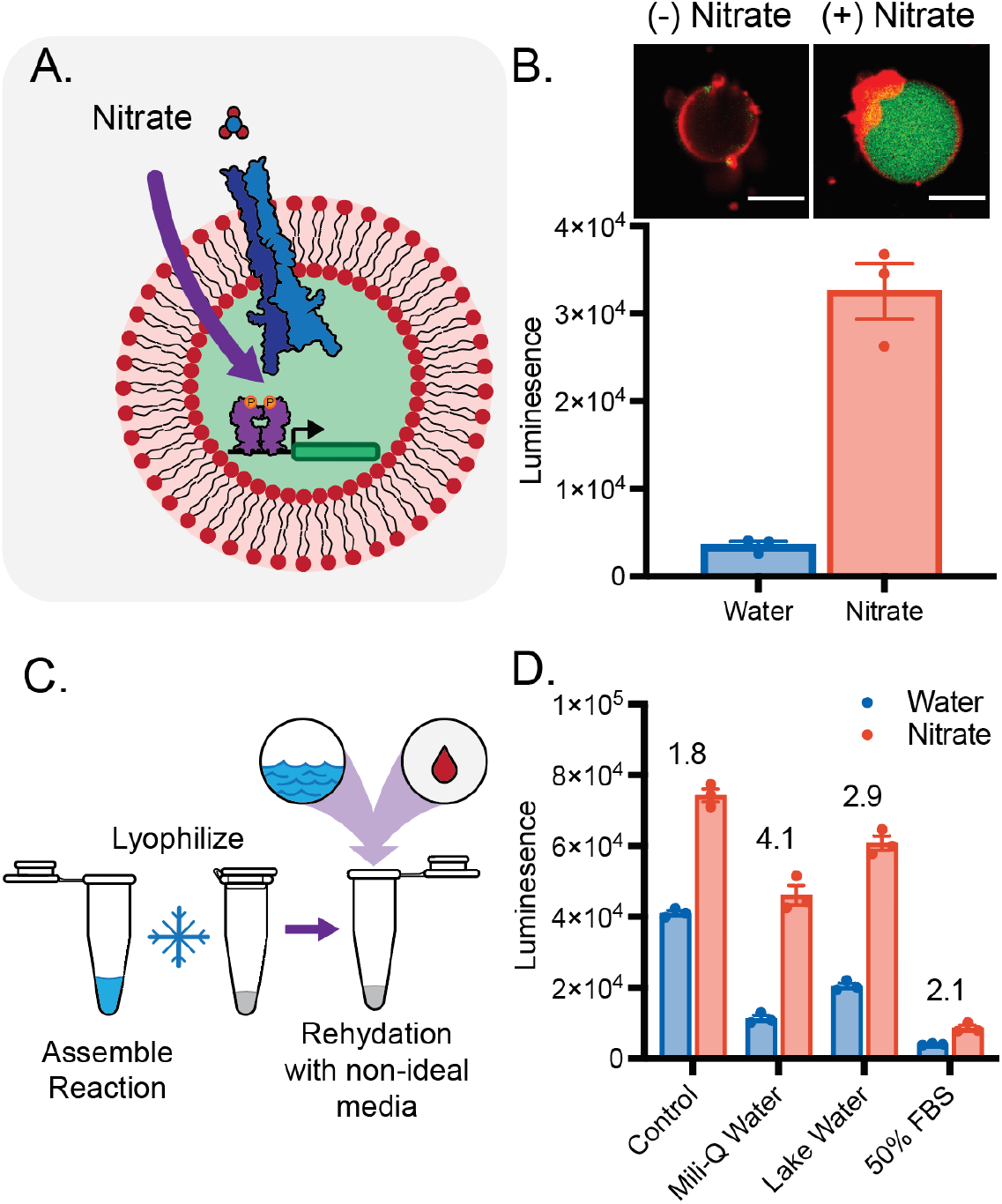
Towards the deployment of two-component sensors for real world sensing. (A) The cell-free expressed NarX-L system can be encapsulated into giant unilamellar vesicles. (B) Signaling and subsequent mEGFP expression can be visualized on the microscope (top). Rhodamine labeled membranes are colored in red and mEGFP are colored in green. Scale bars are 10 µm. Expression of luciferase inside giant unilamellar vesicles can be detected in bulk solution (bottom). (C) Lyophilized reactions remain active, following rehydration with non-ideal media. (D) NarX-L enables higher expression in the presence of 100 µM nitrate compared to the absence of nitrate when rehydrated in Milli-Q water, lake water, and 50% FBS following lyophilization. Fold change in luminescence in response to nitrate is indicated above each condition. All error bars represent the S. E. M. for *n=3* independent replicates.

To demonstrate our system’s ability to be deployed out of the lab to sense ligands in non-ideal matrices, we lyophilized reactions and tested their ability to sense 100 µM nitrate, approximately the EPA limit of nitrate (*30*), when rehydrated with lab-grade Milli-Q water, water from Lake Michigan, and fetal bovine serum (FBS) (Fig. 5C). As a control, we flash froze reactions and allowed them to proceed without lyophilization. We found that lyophilizing vesicles and cell-free components separately produced better results. In reactions with vesicles present during lyophilization, vesicles were found to often float to the top, likely inhibiting membrane-bound kinase expression, and producing low inducible signal (Fig. S10). To remediate this, we lyophilized vesicles and reactions separately. When rehydrating reactions, we rehydrated vesicles with water and then mixed the rehydrated vesicles, lyophilized cell-free components, and rehydration media (water, lake water, or FBS). We then allowed reactions to proceed at 30°C. We found that we could sense 100 µM nitrate in all conditions, however the expression and fold change in luminescence varied in each rehydration media, likely due to different interactions with membranes and transcription/translation machinery (Fig. 5D). Combined, these data demonstrate the ability of cell-free expressed bacterial two-component systems to be integrated and functional in contexts amenable for real-world deployment. We furthermore envision that these systems could be leveraged to engineer targeted drug delivery systems or for point-of-use biosensing.

## Discussion

Here, we show that the bacterial TCS, NarX-L, can be functionally reconstituted into a cell-free system. Leveraging the tunability of cell-free systems, we demonstrate how reaction composition and membrane design may be used to tune the performance of a model two component system, NarX-L. Employing rational protein design we generate new nickel and iron sensors and explore how transmembrane sequences may require membranes with different physiochemical properties for optimal function. Much work has been done characterizing and engineering membrane bound TCS’s in both prokaryotic and eukaryotic systems (*21, 26, 28, 46, 48*); however, we show that these systems can be reconstituted *in vitro* into synthetic membranes which may serve as a platform to both probe biophysical interactions between protein and membrane components and enable new membrane-based biosensors and therapeutics.

Characterizing how membrane-protein interactions affect membrane protein folding and activity is integral to uncovering how such interactions may contribute to disease or be leveraged in biotechnologies (*9, 49, 50*). Toward this goal, *in vitro* systems have been used to characterize purified protein activity and cotranslational folding of membrane proteins. Previous work has explored how membrane composition and mechanical properties affect protein integration and folding into synthetic membranes (*16, 42*). Separately, membrane proteins have been extracted from cells and studied in synthetic membranes. This process allows for the study of specific membrane properties on protein function and has enabled a better understanding of energy generation (*51*), lipid sensing (*40*), and cellular signaling (*39, 48*). However, to the best of our knowledge, we are the first to demonstrate and characterize the folding and activity of a cell-free expressed transmembrane protein-to-genetic reporter signaling pathway. Such a system enables the characterization of protein folding and function in one pot and does not require protein purification and reconstitution. Through this work, we found that membrane viscosity greatly impacts protein insertion and folding. Interestingly, we found that PE and PG lipids allowed for the highest luminescence in response to nitrate, which are the most abundant headgroups found in *E. coli*, NarX’s native membrane. Furthermore, the relationship between membrane properties and activity for our chimeric iron sensor agrees with previously reported *in vivo* data demonstrating that RssA is inhibited by saturated fatty acids (*52*). Together, this data demonstrates that this platform may approximate activity in membranes with similar biophysical properties and could provide a route to assess the relationship between membrane physiochemical properties and membrane protein folding and activity in high-throughput.

Extending this platform beyond biophysical characterization of transmembrane signaling could enable the creation of new membrane-based biosensors and therapeutics. Cell-free systems have been shown to be powerful platforms to build biosensors (*6–8, 53*) and therapeutics (*3, 5*). Over the last 20 years, membranes have been used to encapsulate cell-free systems, towards the goal of engineering an artificial cell (*1, 9*). Such encapsulation systems have been shown to enable prolonged protein expression (*10*), release of cargo in response to environmental cues (*11, 17*), as well as protection from degradation (*12*). However, to date such artificial cellular systems have relied on the nonspecific transport of materials across the membrane to initiate a response. This approach limits the types of signals which can be sensed to those that passively cross the bilayer membrane and when non-specific pores are used to improve molecular transport, they risk leakage of encapsulated components required for function. Together the limited design of membranes that effectively utilize transmembrane proteins has vastly limited the potential applications of membrane-augmented cell-free systems to date.

Biological cells transduce most environmental signals across membranes via transmembrane receptors, which allow cells to sense and respond to signals without moving material across the membrane. The ability to incorporate transmembrane receptors into synthetic membranes will be a powerful tool to rapidly expand the sensing capabilities of cell-mimetic systems. Here, we demonstrate the functional integration of cell-free expressed bacterial transmembrane signaling cascades in synthetic membranes. Our results demonstrate that protein activity is greatly affected by the membrane physiochemical environment and that proteins with different sequences require different lipid environments for proper function. These findings underscore how the lipid-nano environment affects the function of transmembrane proteins and highlight how the membrane environment will be a critical part to consider in the design of systems which leverage transmembrane receptors (*54*). By incorporating genetic parts already developed for bacterial two-component systems (*21, 24, 26*), we believe that transmembrane signaling in response to a wide array of ligands including metals, small molecules, and proteins is within reach. Further, novel receptor systems have been developed in cellular systems to enable cells to sense and respond to new ligands (*55–60*), and their integration into cell-free systems could greatly expand sensing capabilities of cell-mimetic systems. Lipid based particles have proven to be powerful drug delivery vehicles and have been used to deliver cell-free expressed proteins (*47*). Engineering materials capable of sensing and responding to stimuli would combine the responsiveness of cell-based therapeutics with the tunability of cell-free systems, drastically improving our ability to engineer targeted therapeutic delivery systems and new membrane-based materials.

## Materials and Methods Materials

1,2-dimyristoyl-sn-glycero-3-phosphocholine (DMPC), 1-palmitoyl-2-oleoyl-glycero-3-phosphocholine (POPC), 1,2-dioleoyl-sn-glycero-3-phosphocholine (DOPC), 1-palmitoyl-2-oleoyl-sn-glycero-3-phosphoethanolamine (POPE), 1-palmitoyl-2-oleoyl-sn-glycero-3-phospho-(1’-rac-glycerol) (sodium salt) (POPG), E coli Polar Lipid Extract, Cholesterol, and 1,2-dioleoyl-sn-glycero-3-phosphoethanolamine-N-(7-nitro-2-1,3-benzoxadiazol-4-yl)1,2-dioleoyl-snglycero-3-phosphoethanolamine-N-(lissamine rhodamine B sulfonyl) (18:1 Rhodamine) were purchased from Avanti Polar Lipids. Cell-extract (myTxTl) was purchased from Arbor Bio Sciences. gBlocks and primers were ordered from Integrated DNA technologies and DNA was amplified and assembled using enzymes from Thermo Fisher. NanoGlo luciferase was purchased from Promega.

### Plasmid Design and Construction

DNA encoding NarX, NarL, YdfJ promoter, and nanoluciferase were received as gifts from Michael Jewett and were recloned into cell-free backbones using Gibson Assembly. gBlocks encoding kinase chimeras and primers for cloning were ordered from Integrated DNA technologies. Enzymes and buffers required for PCR and cloning were purchased from Thermo Fisher.

### Vesicle Preparation

Vesicles were prepared using the thin film hydration method. Briefly, lipid was deposited into a glass vial and dried with a stream of nitrogen and placed under vacuum for 3 hours. Films were then rehydrated in Milli-Q water for a minimum of 3 hours, and up to overnight. Vesicles were then vortexed and extruded 21x through a 100 nm polycarbonate filter.

### Cell-Free Protein Synthesis Reactions

Protein expression was performed using the myTxTl system (Arbor Biosceinces). 5 µL reactions were assembled with 10 mM lipid, 0.1 nM plasmid encoding T7 RNA polymerase, 9.9 nM plasmids encoding NarX, NarL, and reporter gene (nano luciferase or GFP) in specified ratios, and various concentration of ligands. Cell-free reactions were allowed to progress at 30°C for 16 hours.

Luciferase luminescence was read using the NanoGlo luciferase system and GFP fluorescence (ex. 480 nm, em. 507 nm) was read using a Molecular Devices Spectra Max i3 plate reader. To calculate fold change, luminescence values in the presence of nitrate were divided by the luminescence values in the absence of nitrate for each replicate.

### Western Blotting

1 µL of cell-free reaction was diluted to 15 µL in Laemlli buffer and heated at 95 °C for 10 minutes. Samples were then loaded and run on a 12% Mini-PROTEAN TGX Precast Protein Gel (Bio-Rad) at 150 V for 90 minutes. Wet transfer was performed onto a PVDF membrane (Bio-Rad) for 45 min at 100 V. Membranes were then blocked for an hour at room temperature in 5% milk in TBST (pH 7.6: 50 mM Tris, 150 mM NaCl, HCl to pH 7.6, 0.1% Tween) and incubated for 1 hour at room temperature or overnight at 4 °C with primary solution (anti-Flag (Sigma F1804) or anti-Myc (ab32), diluted 1:1000 in 5% milk in TBST). Primary antibody solution was decanted, and the membrane was washed three times for 5 minutes in TBST and then incubated in secondary solution at room temperature for 1 hour (HRP-anti-Mouse (CST 7076) diluted 1:3000 in 5% milk in TBST). Membranes were then washed in TBST and incubated with Clarity Western ECL Substrate (Bio-Rad) for 5 min. Blots were imaged using an Azure Biosystems c280 imager and band intensities were quantified with ImageJ.

### Lyophilization of reactions

Cell-free reactions were prepared as above, omitting vesicles and ligand. Reactions and vesicles were then flash frozen and lyophilized overnight. Following lyophilization, vesicles were rehydrated in water and cell free reactions were rehydrated in various media, such as Mili-Q water, Lake Water from Lake Michigan, and 50% FBS containing either 100 µM nitrate or an equivalent volume of water. Vesicles and reactions were allowed to rehydrate for 30 minutes on ice. Finally, vesicles were added to each reaction to bring the total volume to 5 µL and reactions were allowed to proceed at 30 °C for 16 hours.

### Encapsulation of TCS into vesicles

Cell-free machinery was encapsulated into vesicles using the water-in-oil emulsion method. A lipid film containing a molar ratio 1 POPC : 2 Cholesterol and 0.1 mol% rhodamine conjugate lipid was prepared. Films were then rehydrated with mineral oil and vortexed until the lipid was resuspended. The aqueous inner phase containing cell-free components and plasmids encoding NarX, NarL, and nanoluciferase or mEGFP were added to the lipid-oil mixture and briefly vortexed to produce an emulsion. Emulsions were then incubated for 5 minutes on ice and layered on top of outer solution (100 mM HEPES, 200 mM Glucose). The layered solutions were incubated on ice for 5 minutes and then centrifuged at 18,000 RCF for 15 minutes at 4 °C. The top oil phase was removed with a pipette and the vesicle pellet was collected and transferred to a new tube containing an equal volume of outer solution. Vesicles were then centrifuged at 12,000 RCF for 5 minutes at 4 °C. The vesicle pellet was then collected and divided into two. Outer solution containing RNAse and 1 mM nitrate was then added to 25 µL and samples were incubated at 30 °C for 16 hours.

For samples containing the luciferase plasmid, 0.5 µL of NanoGlo Luciferase substrate was added to the reaction, and reactions were read on a Molecular Devices Spectra Max i3 plate reader. For reactions containing a downstream mEGFP, 1 µL of the cell-free reaction was added to 100 µL of outer solution in a glass bottom 96 well plate (Corning). Samples were then imaged using a 20x objective on a Nikon confocal microscope. Images were analyzed using the Nikon NIS software.

## Acknowledgments

We thank A. Silverman for helpful discussions regarding the design of cell-free expressed two component systems, and C. Hilburger and the Kamat lab for giving feedback and proof reading of the manuscript. This work was supported in part by the National Science Foundation under Grant No. 1844219, 1844336, 2145050. We thank the financial support of the CBC-NU Cell-free Biomanufacturing Institute funded by the U.S. Army Contracting Command Award W52P1J-21-9-3023 (J.A.P.). J.A.P. gratefully acknowledges support from the NSF Graduate Research Fellowship Program, the Ryan Fellowship, and the International Institute for Nanotechnology at Northwestern University. N.R.G. was supported by a McCormick Summer Research Fellowship Award.

## References

1. A. D. Silverman, A. S. Karim, M. C. Jewett, Cell-free gene expression: an expanded repertoire of applications. Nature Reviews Genetics 2019 21:3. 21, 151–170 (2019).

2. M. W. Nirenberg, J. H. Matthaei, The dependence of cell-free protein synthesis in E. coli upon naturally occurring or synthetic polyribonucleotides. Proc Natl Acad Sci U S A. 47, 1588–1602 (1961).

3. A. C. Hunt, B. Vögeli, W. K. Kightlinger, D. J. Yoesep, A. Krüger, M. C. Jewett, bioRxiv, in press, doi:10.1101/2021.11.04.467378.

4. K. F. Warfel, A. Williams, D. A. Wong, S. E. Sobol, P. Desai, J. Li, Y.-F. Chang, M. P. Delisa, S. Karim, M. C. Jewett, R. Frederick, bioRxiv, in press, doi:10.1101/2022.08.10.503507.

5. J. C. Stark, T. Jaroentomeechai, T. D. Moeller, J. M. Hershewe, K. F. Warfel, B. S. Moricz, M. Martini, R. S. Dubner, K. J. Hsu, T. C. Stevenson, B. D. Jones, M. P. DeLisa, M. C. Jewett, On-demand biomanufacturing of protective conjugate vaccines. Sci Adv. 7 (2021), doi:10.1126/SCIADV.ABE9444/SUPPL_FILE/ABE9444_SM.PDF.

6. W. Thavarajah, A. D. Silverman, M. S. Verosloff, N. Kelley-Loughnane, M. C. Jewett, J. B. Lucks, Point-of-Use Detection of Environmental Fluoride via a Cell-Free Riboswitch-Based Biosensor. ACS Synth Biol. 9, 10–18 (2020).

7. J. K. Jung, K. K. Alam, M. S. Verosloff, D. A. Capdevila, M. Desmau, P. R. Clauer, J. W. Lee, P. Q. Nguyen, P. A. Pastén, S. J. Matiasek, J. F. Gaillard, D. P. Giedroc, J. J. Collins, J. B. Lucks, Cell-free biosensors for rapid detection of water contaminants. Nature Biotechnology 2020 38:12. 38, 1451–1459 (2020).

8. P. Q. Nguyen, L. R. Soenksen, N. M. Donghia, N. M. Angenent-Mari, H. de Puig, A. Huang, R. Lee, S. Slomovic, T. Galbersanini, G. Lansberry, H. M. Sallum, E. M. Zhao, J. B. Niemi, J. J. Collins, Wearable materials with embedded synthetic biology sensors for biomolecule detection. Nature Biotechnology 2021 39:11. 39, 1366–1374 (2021).

9. N. S. Kruyer, W. Sugianto, B. I. Tickman, D. Alba Burbano, V. Noireaux, J. M. Carothers, P. Peralta-Yahya, Membrane Augmented Cell-Free Systems: A New Frontier in Biotechnology. ACS Synth Biol. 10, 670–681 (2021).

10. V. Noireaux, A. Libchaber, A vesicle bioreactor as a step toward an artificial cell assembly. Proc Natl Acad Sci U S A. 101, 17669–17674 (2004).

11. K. P. Adamala, D. A. Martin-Alarcon, K. R. Guthrie-Honea, E. S. Boyden, Engineering genetic circuit interactions within and between synthetic minimal cells. Nature Chemistry 2016 9:5. 9, 431–439 (2016).

12. M. A. Boyd, W. Thavarajah, J. B. Lucks, N. P. Kamat, bioRxiv, in press, doi:10.1101/2022.03.02.482665.

13. J. M. Heili, K. Stokes, N. J. Gaut, C. Deich, J. Gomez-Garcia, B. Cash, M. R. Pawlak, A. E. Engelhart, K. P. Adamala, bioRxiv, in press, doi:10.1101/2022.01.03.474826.

14. J. A. Schoborg, J. M. Hershewe, J. C. Stark, W. Kightlinger, J. E. Kath, T. Jaroentomeechai Natarajan, M. P. DeLisa, M. C. Jewett, A cell-free platform for rapid synthesis and testing of active oligosaccharyltransferases. Biotechnol Bioeng. 115, 739–750 (2018).

15. D. Blanken, D. Foschepoth, A. C. Serrão, C. Danelon, Genetically controlled membrane synthesis in liposomes. Nature Communications 2020 11:1. 11, 1–13 (2020).

16. M. L. Jacobs, M. A. Boyd, N. P. Kamat, Diblock copolymers enhance folding of a mechanosensitive membrane protein during cell-free expression. Proc Natl Acad Sci U S A. 116, 4031–4036 (2019).

17. D. Toparlak, J. Zasso, S. Bridi, M. D. Serra, P. MacChi, L. Conti, M. L. Baudet, S. S. Mansy, Artificial cells drive neural differentiation. Sci Adv. 6, 4920–4938 (2020).

18. I. Gispert, J. W. Hindley, C. P. Pilkington, H. Shree, L. M. C. Barter, O. Ces, Y. Elani, Stimuli-responsive vesicles as distributed artificial organelles for bacterial activation. Proceedings of the National Academy of Sciences. 119, e2206563119 (2022).

19. J. W. Hindley, D. G. Zheleva, Y. Elani, K. Charalambous, L. M. C. Barter, P. J. Booth, C. L. Bevan, R. v. Law, O. Ces, Building a synthetic mechanosensitive signaling pathway in compartmentalized artificial cells. Proc Natl Acad Sci U S A. 116, 16711–16716 (2019).

20. F. Jacob-Dubuisson, A. Mechaly, J. M. Betton, R. Antoine, Structural insights into the signalling mechanisms of two-component systems. Nature Reviews Microbiology 2018 16:10. 16, 585–593 (2018).

21. J. T. Lazar, J. J. Tabor, Bacterial two-component systems as sensors for synthetic biology applications. Curr Opin Syst Biol. 28, 100398 (2021).

22. P. Zhang, J. Yang, E. Cho, Y. Lu, Bringing Light into Cell-Free Expression. ACS Synth Biol. 9, 2144–2153 (2020).

23. J. Yuan, F. Jin, T. Glatter, V. Sourjik, Osmosensing by the bacterial PhoQ/PhoP two-component system. Proc Natl Acad Sci U S A. 114, E10792–E10798 (2017).

24. B. P. Landry, R. Palanki, N. Dyulgyarov, L. A. Hartsough, J. J. Tabor, Phosphatase activity tunes two-component system sensor detection threshold. Nature Communications 2018 9:1. 9, 1–10 (2018).

25. H. Liang, X. Deng, M. Bosscher, Q. Ji, M. P. Jensen, C. He, Engineering bacterial two-component system PmrA/PmrB to sense lanthanide ions. J Am Chem Soc. 135, 2037–2039 (2013).

26. S. R. Schmidl, F. Ekness, K. Sofjan, K. N. M. Daeffler, K. R. Brink, B. P. Landry, K. P. Gerhardt, N. Dyulgyarov, R. U. Sheth, J. J. Tabor, Rewiring bacterial two-component systems by modular DNA-binding domain swapping. Nature Chemical Biology 2019 15:7. 15, 690–698 (2019).

27. W. R. Whitaker, S. A. Davis, A. P. Arkin, J. E. Dueber, Engineering robust control of two-component system phosphotransfer using modular scaffolds. Proc Natl Acad Sci U S A. 109, 18090–18095 (2012).

28. A. Mazé, Y. Benenson, Artificial signaling in mammalian cells enabled by prokaryotic two-component system. Nature Chemical Biology 2019 16:2. 16, 179–187 (2019).

29. J. Hansen, E. Mail, K. K. Swaminathan, J. Schreiber, B. Angelici, Y. Benenson, Transplantation of prokaryotic two-component signaling pathways into mammalian cells. Proc Natl Acad Sci U S A. 111, 15705–15710 (2014).

30. Estimated Nitrate Concentrations in Groundwater Used for Drinking | US EPA, (available at https://www.epa.gov/nutrient-policy-data/estimated-nitrate-concentrations-groundwater-used-drinking).

31. R. Marshall, V. Noireaux, Synthetic biology with an all E. coli TXTL system: Quantitative characterization of regulatory elements and gene circuits. Methods in Molecular Biology. 1772, 61–93 (2018).

32. M. Serizawa, J. Sekiguchi, The Bacillus subtilis YdfHI two-component system regulates the transcription of ydfJ, a member of the RND superfamily. Microbiology (N Y). 151, 1769–1778 (2005).

33. J. M. Hershewe, K. F. Warfel, S. M. Iyer, J. A. Peruzzi, C. J. Sullivan, E. W. Roth, M. P. DeLisa, N. P. Kamat, M. C. Jewett, Improving cell-free glycoprotein synthesis by characterizing and enriching native membrane vesicles. Nature Communications 2021 12:1. 12, 1–12 (2021).

34. J. J. Wuu, J. R. Swartz, High yield cell-free production of integral membrane proteins without refolding or detergents. Biochimica et Biophysica Acta (BBA) - Biomembranes. 1778, 1237–1250 (2008).

35. M. C. Jewett, K. A. Calhoun, A. Voloshin, J. J. Wuu, J. R. Swartz, An integrated cell-free metabolic platform for protein production and synthetic biology. Mol Syst Biol. 4, 220 (2008).

36. A. D. Silverman, N. Kelley-Loughnane, J. B. Lucks, M. C. Jewett, Deconstructing Cell-Free Extract Preparation for in Vitro Activation of Transcriptional Genetic Circuitry. ACS Synth Biol. 8, 403–414 (2019).

37. C. E. Hilburger, M. L. Jacobs, K. R. Lewis, J. A. Peruzzi, N. P. Kamat, Controlling Secretion in Artificial Cells with a Membrane and Gate. ACS Synth Biol. 8, 1224–1230 (2019).

38. M. L. Jacobs, J. Steinkühler, A. Lemley, M. J. Larmore, L. C. T. Ng, S. M. Cologna, P. G. DeCaen, N. P. Kamat, bioRxiv, in press, doi:10.1101/2022.05.20.492859.

39. M. G. Gutierrez, K. S. Mansfield, N. Malmstadt, The Functional Activity of the Human Serotonin 5-HT1A Receptor Is Controlled by Lipid Bilayer Composition. Biophys J. 110, 2486–2495 (2016).

40. S. Ballweg, E. Sezgin, M. Doktorova, R. Covino, J. Reinhard, D. Wunnicke, I. Hänelt, I. Levental, G. Hummer, R. Ernst, Regulation of lipid saturation without sensing membrane fluidity. Nature Communications 2020 11:1. 11, 1–13 (2020).

41. J. A. Peruzzi, J. Steinkühler, T. Q. Vu, T. F. Gunnels, P. Lu, D. Baker, N. P. Kamat, bioRxiv, in press, doi:10.1101/2022.06.01.494374.

42. N. J. Harris, E. Reading, K. Ataka, L. Grzegorzewski, K. Charalambous, X. Liu, R. Schlesinger, J. Heberle, P. J. Booth, Structure formation during translocon-unassisted co-translational membrane protein folding. Scientific Reports 2017 7:1. 7, 1–15 (2017).

43. M. R. Sanders, H. E. Findlay, P. J. Booth, Lipid bilayer composition modulates the unfolding free energy of a knotted α-helical membrane protein. Proc Natl Acad Sci U S A. 115, E1709–E1808 (2018).

44. B. Mensa, N. F. Polizzi, K. S. Molnar, A. M. Natale, T. Lemmin, W. F. Degrado, Allosteric mechanism of signal transduction in the two-component system histidine kinase PhoQ. Elife. 10 (2021), doi:10.7554/ELIFE.73336.

45. K. Pluhackova, A. Horner, Native-like membrane models of E. coli polar lipid extract shed light on the importance of lipid composition complexity. BMC Biology 2021 19:1. 19, 1–22 (2021).

46. R. Utsumi, R. E. Brissette, A. Rampersaud, S. A. Forst, K. Oosawa, M. Inouye, Activation of Bacterial Porin Gene Expression by a Chimeric Signal Transducer in Response to Aspartate. Science (1979). 245, 1246–1249 (1989).

47. N. Krinsky, M. Kaduri, A. Zinger, J. Shainsky-Roitman, M. Goldfeder, I. Benhar, D. Hershkovitz, A. Schroeder, Synthetic Cells Synthesize Therapeutic Proteins inside Tumors. Adv Healthc Mater. 7, 1701163 (2018).

48. M. E. Inda, M. Vandenbranden, A. Fernández, D. de Mendoza, J. M. Ruysschaert, L. E. Cybulski, A lipid-mediated conformational switch modulates the thermosensing activity of DesK. Proc Natl Acad Sci U S A. 111, 3579–3584 (2014).

49. I. Levental, E. Lyman, Regulation of membrane protein structure and function by their lipid nano-environment. Nature Reviews Molecular Cell Biology 2022, 1–16 (2022).

50. J. T. Marinko, H. Huang, W. D. Penn, J. A. Capra, J. P. Schlebach, C. R. Sanders, Folding and Misfolding of Human Membrane Proteins in Health and Disease: From Single Molecules to Cellular Proteostasis. Chem Rev. 119, 5537–5606 (2019).

51. S. Berhanu, T. Ueda, Y. Kuruma, Artificial photosynthetic cell producing energy for protein synthesis. Nature Communications 2019 10:1. 10, 1–10 (2019).

52. H. C. Lai, P. C. Soo, J. R. Wei, W. C. Yi, S. J. Liaw, Y. T. Horng, S. M. Lin, S. W. Ho, S. Swift, P. Williams, The RssAB two-component signal transduction system in Serratia marcescens regulates swarming motility and cell envelope architecture in response to exogenous saturated fatty acids. J Bacteriol. 187, 3407–3414 (2005).

53. A. D. Silverman, U. Akova, K. K. Alam, M. C. Jewett, J. B. Lucks, Design and Optimization of a Cell-Free Atrazine Biosensor. ACS Synth Biol. 9, 671–677 (2020).

54. I. Levental, E. Lyman, Regulation of membrane protein structure and function by their lipid nano-environment. Nature Reviews Molecular Cell Biology 2022, 1–16 (2022).

55. H. J. Jackson, S. Rafiq, R. J. Brentjens, Driving CAR T-cells forward. Nat Rev Clin Oncol. 13 (2016), pp. 370–383.

56. S. H. Park, A. Zarrinpar, W. A. Lim, Rewiring MAP kinase pathways using alternative scaffold assembly mechanisms. Science (1979). 299, 1061–1064 (2003).

57. J. Hansen, Y. Benenson, Synthetic biology of cell signaling. Nat Comput. 15, 5–13 (2016).

58. Y. Benenson, Biomolecular computing systems: Principles, progress and potential. Nat Rev Genet. 13, 455–468 (2012).

59. A. Mazé, Y. Benenson, Artificial signaling in mammalian cells enabled by prokaryotic two-component system. Nat Chem Biol. 16, 179–187 (2020).

60. T. Strittmatter, Y. Wang, A. Bertschi, L. Scheller, P. C. Freitag, P. G. Ray, P. Stuecheli, J. v. Schaefer, T. Reinberg, D. Tsakiris, A. Plückthun, H. Ye, M. Fussenegger, Programmable DARPin-based receptors for the detection of thrombotic markers. Nature Chemical Biology 2022, 1–10 (2022).

